# Boosting carbon fixation and microbial dynamics in the coastal sediment ecosystem through large-scale cultivation of *Gracilariopsis lemaneiformis*

**DOI:** 10.64898/2026.07.01.735803

**Authors:** Pengbing Pei, Yinglong Chen, Muhammad Aslam, Chuanyu Wu, Weihui Zeng, Hong Du

**Author notes:** **Corresponding author:** Hong Du. Both authors contributed equally to the article (Pengbing Pei, Yinglong Chen).

## Abstract

Microorganisms are the key drivers of carbon cycling in coastal marine sediment ecosystems, significantly influencing carbon storage and release during *Gracilariopsis lemaneiformis* cultivation. This study employed 16S rRNA sequencing, a high-throughput qPCR chip, and carbon isotope labeling to assess the impact of *G. lemaneiformis* cultivation on carbon cycling processes in coastal sediments. A comparative analysis was conducted between cultivated zones (GZ) of *G. lemaneiformis* and adjacent control zones (CZ). The results indicated that macroalgae cultivation significantly modified sediment-seawater exchange dynamics and accelerated carbon cycling within coastal marine sediment ecosystems. Furthermore, *G. lemaneiformis* cultivation increased the abundance of genes linked to polysaccharide degradation and carbon fixation pathways, thereby enhancing carbon cycling efficiency. The ecosystem multifunctional index, calculated based on carbon fixation gene abundance, was significantly higher in GZ compared to CZ. Incubation experiments using ^13^C-NaHCO_3_ demonstrated that cultivation markedly elevated the carbon fixation rate of sediment, emphasizing a higher potential for carbon sequestration in sedimentary environments cultivated with macroalgae. Additionally, cultivation significantly altered sediment microbial communities, simplifying their structural complexity. Key microbial taxa identified via k-core species analysis—including Subgroup10 of Desulfobacterota and MBNT15, correlated strongly with carbon fixation rates, indicating their pivotal roles in sediment carbon cycling processes. This study provides critical insights into how large-scale macroalgae cultivation influences coastal carbon dynamics and informs strategies for optimizing carbon management in aquaculture ecosystems.

**Importance:** *Gracilariopsis lemaneiformis* cultivation is a widely promoted eco-aquaculture model along Southeast China’s coast. This study finds it can drastically boost carbon sequestration efficiency in coastal sediments. Through field comparison and isotope tracing, we confirm that key carbon-fixing microbes in farmed sediments show significantly higher activity, with far stronger carbon storage potential than non-farmed zones. This research delivers direct scientific evidence for scaling up macroalgae farming to unlock new blue carbon gains, and offers a practical new solution for optimizing carbon management in aquaculture ecosystems.

## 1 Introduction

Human activities have increased greenhouse gas emissions in coastal areas, eroding marine ecosystems and disrupting their dynamic equilibrium (Doney et al., 2012; IPCC, 2023). Climate change mitigation entails two key strategies: reducing emissions from fossil fuels and industry, and enhancing carbon sinks through innovations spanning from carbon capture technologies to nature-based approaches like sustainable agriculture and ecological restoration (Bui et al., 2018; Wang et al., 2022; IPCC, 2023; Wang et al., 2025). As a pivotal carbon sink within the Earth’s system, the ocean has garnered growing recognition for its immense carbon sequestration potential (Koeve et al., 2024). This process is largely driven by blue carbon ecosystems, specifically mangroves, seagrass beds, and salt marshes, which absorb and sequester atmospheric CO_2_ through photosynthesis (McLeod et al., 2011). Despite occupying less than 0.5% of the seafloor area, these coastal ecosystems account for over 50% of the carbon storage in marine sediments (Macreadie et al., 2021).

Emerging as a promising addition to these natural carbon sink systems, large-scale seaweed aquaculture represents a proactive strategy to potentially enhance the ocean’s carbon sequestration capacity (Wang et al., 2025). A significant portion of the organic carbon produced during macroalgal photosynthesis contributes to long-term carbon sequestration through processes such as sediment burial, deep-sea transport, and transformation into refractory organic carbon (Gao et al., 2022). Beyond the photosynthetic activity of macroalgae itself, microbial communities in aquaculture areas also play a critical role in carbon sequestration via biogeochemical cycling (Chen et al., 2025; Sun et al., 2023). Microorganisms are pivotal drivers of biogeochemical cycles, with bacteria playing a predominant role in the carbon and nitrogen cycles within various ecosystems (Boyd et al., 2010; Crowther et al., 2019; Pei et al., 2024a & 2024b). Previous research has focused on the composition of microbial communities associated with macroalgae (Egan et al., 2012; Xie et al., 2017; Wang et al., 2020; Pei et al., 2021; Xu et al., 2022). Meanwhile, microorganisms within aquaculture ecosystem significantly influence the carbon sequestration process, modulating the transformation and storage of organic carbon (Mitra et al., 2014; Zhang et al., 2023). These microorganisms expedite the biological cycle of the bio-pump mechanism (Huang et al., 2024), while facilitating the decomposition and dissolution of organic carbon via the microbial carbon pump process. This dual action promotes the formation of recalcitrant organic carbon, enhancing marine carbon storage capacity through sequestration (Jiao et al., 2010; Gao et al., 2022). Recent studies have begun exploring how macroalgae cultivation influences microbial community biogeochemical cycles and their underlying mechanisms. For instance, kelp cultivation has been shown to significantly alter microbial community diversity and network structure, amplifying biogeochemical cycling functions within these ecosystems (Sun et al., 2023). Enhanced biogeochemical cycling in aquaculture areas correlates with increased bacterioplankton richness and intensified interspecies interactions (Choi et al., 2016; Selvarajan et al., 2019; Xie et al., 2024). This pattern is consistent across diverse coastal ecosystems (Miranda et al., 2022; Qian et al., 2023). However, the complex interactions between microbial communities and carbon biogeochemical cycles in macroalgae aquaculture ecosystems remain poorly understood, representing a critical gap in current scientific knowledge.

The presence and abundance of key functional genes encoding enzymes are indicative of the carbon cycle potential across diverse habitats (Sun et al., 2023). So far, seven distinct pathways for microbial carbon fixation have been identified (Berg, 2011). Among these, the Calvin-Benson-Bassham (CBB) cycle and the 3-hydroxypropionic acid/4-hydroxybutyric acid (3-HP/4-HB) cycle represent the most significant chemoautotrophic CO₂ fixation pathways in coastal ecosystems (Liu et al., 2022; Mustafa et al., 2024). Relies on the enzyme ribulose-1,5-bisphosphate carboxylase/oxygenase (RubisCO), which is critical for CO₂ fixation (Sporre et al., 2023). In the 3-HP/4-HB cycle, acetyl-CoA carboxylase (ACCase) encoded by the marker gene *accA*-catalyzes the conversion of acetyl-CoA and bicarbonate into malonyl-CoA and acts as a diagnostic marker for this pathway in autotrophic archaea (Pearson et al., 2019; Ding et al., 2021).

Coastal sediment ecosystems have significant potential for carbon sequestration and play a crucial role in the global carbon cycle (Queiros et al., 2019). Chemoautotrophic carbon fixation in coastal sediments, which occupy 7% of the ocean’s area, is estimated at approximately 175 Tg C year⁻¹, representing 22.7% of total oceanic chemoautotrophic carbon fixation (Dai et al., 2022). The carbon cycle in coastal sediments operates as an integrated system, where reductive and oxidative processes closely link carbon, nitrogen, sulfur, hydrogen, and metal cycles, generating substrates that fuel chemoautotrophy (Hu et al., 2024). Anthropogenic inputs of organic matter into estuarine and coastal ecosystems further enhance the oxidation of reduced inorganic nitrogen compounds. This process provides a critical energy source for chemoautotrophic microorganisms, amplifying their role in carbon sequestration (Galloway et al., 2024). Large-scale macroalgae cultivation results in significant deposition of macroalgal detritus into coastal sediments (Feng et al., 2022). The input of organic matter significantly impacts carbon sequestration within coastal sediment ecosystems. Owing to this abundant organic matter input, mangroves, intertidal zones, and estuaries represent key ecosystems frequently studied in scientific research. The influence of environmental factors, including temperature, salinity, and nutrient availability,has been extensively examined (Liu et al., 2022; Hu et al., 2024). Microbial keystone species and their interactions are pivotal drivers of biogeochemical cycling processes (Lin et al., 2023). Large-scale macroalgae cultivation has gained recognition as a nature-based solution for advancing a carbon-negative economy and fostering ecological restoration (Wang et al., 2025).

While the sequestration and transformation of organic matter in coastal sediments have been extensively studied, the role of microbial communities in carbon fixation within aquaculture ecosystems remains understudied. To date, a critical knowledge gap persists regarding how large-scale macroalgae cultivation influences carbon fixation processes in coastal marine sediment ecosystems. We therefore propose the following hypotheses: (1) Large-scale macroalgae cultivation accelerates carbon fixation processes within coastal sediment ecosystems. (2) This acceleration induces a carbon-negative effect in cultivation area sediments, mediated by the dominance of specific microbial communities. To test these hypotheses, paired sediment samples were collected from a macroalgae cultivation area and an adjacent non-cultivation control site. Bacterial communities were analyzed via 16S rRNA gene high-throughput sequencing, and functional genes associated with microbial carbon fixation were quantified using a high-throughput quantitative PCR (HT-qPCR) chip. Carbon fixation rates in sediments were measured through incubation experiments using ^13^C-NaHCO_3_ as the sole carbon substrate. This study provides novel insights into the effect of large-scale macroalgae cultivation on coastal carbon dynamics and supports the formulation of effective carbon management strategies for sustainable aquaculture ecosystems.

## 2 Materials and methods

### 2.1 Site description and sample collection

This study was conducted in January 2024 in the coastal area of Zoumapu Village, Nan’ao Island, China **(Fig. S1a)**, where large-scale cultivation of *Gracilariopsis lemaneiformis* has been practiced for decades. *Gracilariopsis* species have ranked as the second most cultivated macroalgae in China, due to their economic value and ecological restoration potential (Wang et al., 2025).

Six sampling sites were selected from each of two zones i.e., GZ (the *G. lemaneiformis* cultivation zone) and CZ (Control Zone) for this study **(Fig. S1, Tab. S1)**. The control zone area is located in the adjacent sea area, with similar hydrological conditions and no aquaculture activities or other human induced pollution. Surface sediment samples (0.2 m × 0.2 m area, 0-0.1 m depth) were collected using a Peterson grab sampler (Gu et al., 2013; Xie et al., 2017), placed in sterile polyethylene boxes, and immediately stored at 4 °C. A total of 12 samples were collected for incubation experiments, physicochemical analyses, and molecular studies. Samples were transported to the laboratory within 1-3 h. A fraction of each sample for DNA extraction was stored at -80 °C, while the remaining portion was kept at 4 °C for physicochemical characterization and carbon fixation rate measurements.

### 2.2 Measurements of physicochemical parameters

Temperature, salinity, pH, dissolved oxygen (DO), total dissolved solids (TDS), water depth, and coordinates were measured in the overlying seawater column from the in-situ data using an Aqua TROLL® 400 Multiparameter Probe. Water content of the sediment was determined by drying the sediment in a GZX-9070MBE electric-heat forced-draft drying oven (Boxun, China) at 105 °C to constant weight. Results were calculated as dry-to-wet weight ratio. Total carbon (TC), total nitrogen (TN), total sulfur (TS) in sediments were quantified using a Vario EL Cube elemental analyzer (Elementar, Germany). Inorganic nitrogen (NH₄⁺, NO₂⁻, NO₃⁻) was extracted from 5 g of fresh sediment with 2 M KCl and analyzed using UV2400 spectrophotometer (Sunny Hengping, China) following Chinese Standard HJ 634-2012. C/N ratio was calculated as TC divided by TN. TOC (total organic carbon), DOC (dissolved organic carbon), and DIC (dissolved inorganic carbon) in sediments were analyzed using an Elab-TOC/DT analyzer (Elab, China) at the Guangzhou Institute of Chemistry, Chinese Academy of Sciences Co., Ltd., following established protocols (Liu et al., 2022; Bolan et al., 1996; Sheng et al., 2015).

### 2.3 Measurement of carbon cycling genes

A quantitative microbial element cycling (QMEC) qPCR chip, containing primers targeting microbial functional genes involved in carbon fixation, was used to assess the carbon fixation potential of each sample (Zheng et al., 2018). DNA from each sample served as the template for QMEC analysis, quantified via HT-qPCR using a SmartChip Real-Time PCR system (WaferGen Biosystems, Fremont, CA, USA). The bacterial 16S rRNA gene (27F:5’-AGAGTTTGATCMTGGCTCAG-3’/1492R:5’-TACGGYTACCTTGTTACGACTT-3’) was amplified as a reference for normalization (Sun et al., 2023). Reactions were performed in triplicate following the protocol of Zheng et al. (2018). Data quality control was implemented using SmartChip qPCR software, excluding results with non-specific amplification (multiple melting peaks) or amplification efficiencies outside the 80–120% range. Amplification products with a threshold cycle (CT) ≤31 were retained for downstream analysis. Relative gene abundance was calculated as the ratio of each functional gene’s abundance to that of the 16S rRNA gene, following the method of Looft et al. (2012). All QMEC chip-based assays were conducted by Guangzhou Magigene Co. Ltd. (Guangzhou, China) (Looft et al., 2012).

### 2.4 DNA extraction and 16S rDNA gene sequencing

In this study, the amplification of the 16S rRNA gene was performed using universal primers. To increase sequencing throughput, a 16-bp specific barcode sequence was added to the 5’ end of both the forward and reverse primers (27F: 5’-AGAGTTTGATCMTGGCTCAG-3’/1492R:5’-TACGGYTACCTTGTTACGACTT-3’), allowing accurate allocation of sequences to the corresponding samples based on barcode information after sequencing. Third-generation sequencing technology, specifically the PacBio platform, was employed for sequencing analysis. The PCR products were assessed by 1% agarose gel electrophoresis, followed by purification, library construction, and loading for sequencing. All samples in this study were sent to Guangzhou Magigene Co. Ltd. (Guangzhou, China) for sequencing. The sequence data reported in this study have been deposited in the NCBI GenBank database under the accession numbers SRR36222627-SRR36222638. All SRA data are available at https://www.ncbi.nlm.nih.gov/bioproject/PRJNA1370532.

### 2.5 Measurement of sediment carbon fixation rates

Carbon fixation rates (CFR) were measured using the ^13^C-labeled bicarbonate method (Boschker et al., 2014; Sohm et al., 2020). Briefly, sediment slurries were prepared by mixing fresh sediment, sterile-filtered (0.22 μm) seawater, and 60 mM ^13^C-labeled bicarbonate (CAS: 87081-58-1, Aladdin, China) at a ratio of 1:28:1 (g : mL : mL) in penicillin vials. No-treatment controls were prepared for each sample by substituting ^13^C-labeled bicarbonate with sterile-filtered (0.22 μm) water. Vials were flushed with nitrogen gas for 5 minutes to create an anaerobic environment and subsequently sealed. Slurries were incubated in darkness at the in situ overlying seawater temperature in an incubator (Yiheng, China). Incubations were terminated by adding formaldehyde (final concentration 2%). Dead controls were prepared by adding formaldehyde (final concentration 2%) before introducing ^13^C-labeled bicarbonate. Subsequently, 200 μL of 37% hydrochloric acid was added to the slurries, followed by thorough mixing and aeration in a fume hood to remove residual ^13^C-labeled bicarbonate. The slurries were centrifuged to collect the pellet, which was then resuspended in 30 mL of sterile-filtered (0.22 μm) water, and vortexed thoroughly. This centrifugation, resuspension process was repeated three times to yield the ^13^C-labeled bicarbonate, incubated sediment product.

All ^13^C-labeled sediment products and untreated samples were sent to the Third Institute of Oceanography, Ministry of Natural Resources, China, for analysis of ^13^C-organic matter content. The ^13^C-labeled organic matter was quantified using elemental analyzer-isotope ratio mass spectrometry (EA-IRMS, Sercon Integra2, Sercon, UK), following standard protocols for carbon isotope analysis (Zhang et al., 2012; Hayes, 1993). Additional details on sediment CFR measurements are provided in the Supplementary Material (**Fig. S2**).

### 2.6 Statistical analysis

Differences in environmental parameters between GZ and CZ were assessed using one-way ANOVA in IBM SPSS v22. Pearson correlation analysis was conducted to evaluate relationships between environmental parameters. Visualizations were generated using Origin v2024b and Adobe Illustrator v2022. Non-metric multidimensional scaling (n-MDS) and analysis of similarity (ANOSIM), based on Bray-Curtis dissimilarity, were performed to examine differences in carbon fixation cycle gene profiles between the two zones using the “vegan” package in R. The average fold change in the relative abundance of carbon cycle genes between GZ and CZ samples was calculated, with differences evaluated using Student’s t-test. Sediment multifunctionality was quantified based on the relative abundance of biogeochemical cycling genes, following Sun et al. (2023). Differences in the multifunctionality index between GZ and CZ samples were assessed using the Kruskal-Wallis test in IBM SPSS v22.

Alpha diversity indices of bacterial communities were calculated using the “vegan” package. Differences in alpha diversity among samples were compared using ANOVA with Tukey’s HSD test. Principal coordinates analysis (PCoA) based on beta diversity distances was performed using the “vegan” package. Linear discriminant analysis effect size (LEfSe) was conducted on the Biozeron Cloud Platform (http://www.cloud.biomicroclass.com/CloudPlatform) with default parameters (LDA > 2) to identify biomarkers at various taxonomic levels.

Ecological network analysis, including network construction, topological property assessment, module separation, and keystone taxa identification, was performed using the Molecular Ecological Network Analyses (MENA) Pipeline (http://ieg4.rccc.ou.edu/MENA; Deng et al., 2012). Only operational taxonomic units (OTUs) detected in all six samples of each zone were used for network construction, based on random matrix theory (RMT). Networks were visualized using Gephi v0.9.2 (Cherven, 2013). Subnetworks were constructed using the k-core algorithm to identify keystone species in Gephi v0.9.2. Metacoder tree figures and statistical analyses were generated using the Biozeron Cloud Platform. Sankey diagrams illustrating the species composition of keystone and dominant species in GZ and CZ were created using Wekemo Bioincloud (https://www.bioincloud.tech). Mantel tests and network analyses exploring relationships among key species, sediment carbon fixation rates, carbon fixation gene abundance, and environmental parameters were also performed on Wekemo Bioincloud.

## 3 Results

### 3.1 Site physicochemical parameters

The physicochemical parameters of the sampling sites are presented in **Tab. S1**. Sediment total carbon (TC) content exhibited significant spatial heterogeneity (one-way ANOVA, *p* < 0.05), ranging from 6,621 to 12,132 mg kg⁻¹ in GZ and 5,424 to 6,297 mg kg⁻¹ in CZ **(Fig. S3a)**. Sediment ammonium nitrogen (NH₄-N) content was significantly lower in CZ (0.1405–0.2002 mg kg⁻¹) than in GZ (0.1768–0.3088 mg kg⁻¹; one-way ANOVA, *p* < 0.05; **Fig. S3b**).

Pearson’s correlation analysis revealed relationships among environmental factors **(Fig. 1)**. In both GZ and CZ, overlying seawater pH was positively correlated with total dissolved solids (TDS) and salinity (*p* < 0.05). Sediment nitrite nitrogen (NO₂-N) was positively correlated with dissolved organic carbon (DOC) in GZ (*p* < 0.05), with a similar but non-significant trend in CZ (*p* = 0.55). Sediment total sulfur (TS) was negatively correlated with the carbon-to-nitrogen (C/N) ratio in CZ (*p* < 0.05), with a similar but non-significant trend in GZ (*r* = -0.51, *p* = 0.3). Overlying seawater pH was negatively correlated with sediment TS and TC in GZ (*p* < 0.05), but showed an opposite trend in CZ. Overlying seawater relative dissolved oxygen (RDO) was negatively correlated with NH₄-N in GZ (*p* < 0.01), but showed an opposite trend in CZ. In CZ, RDO was negatively correlated with dissolved inorganic carbon (DIC) and DOC (*p* < 0.05), whereas no significant correlations were observed in GZ.

**Figure 1.**
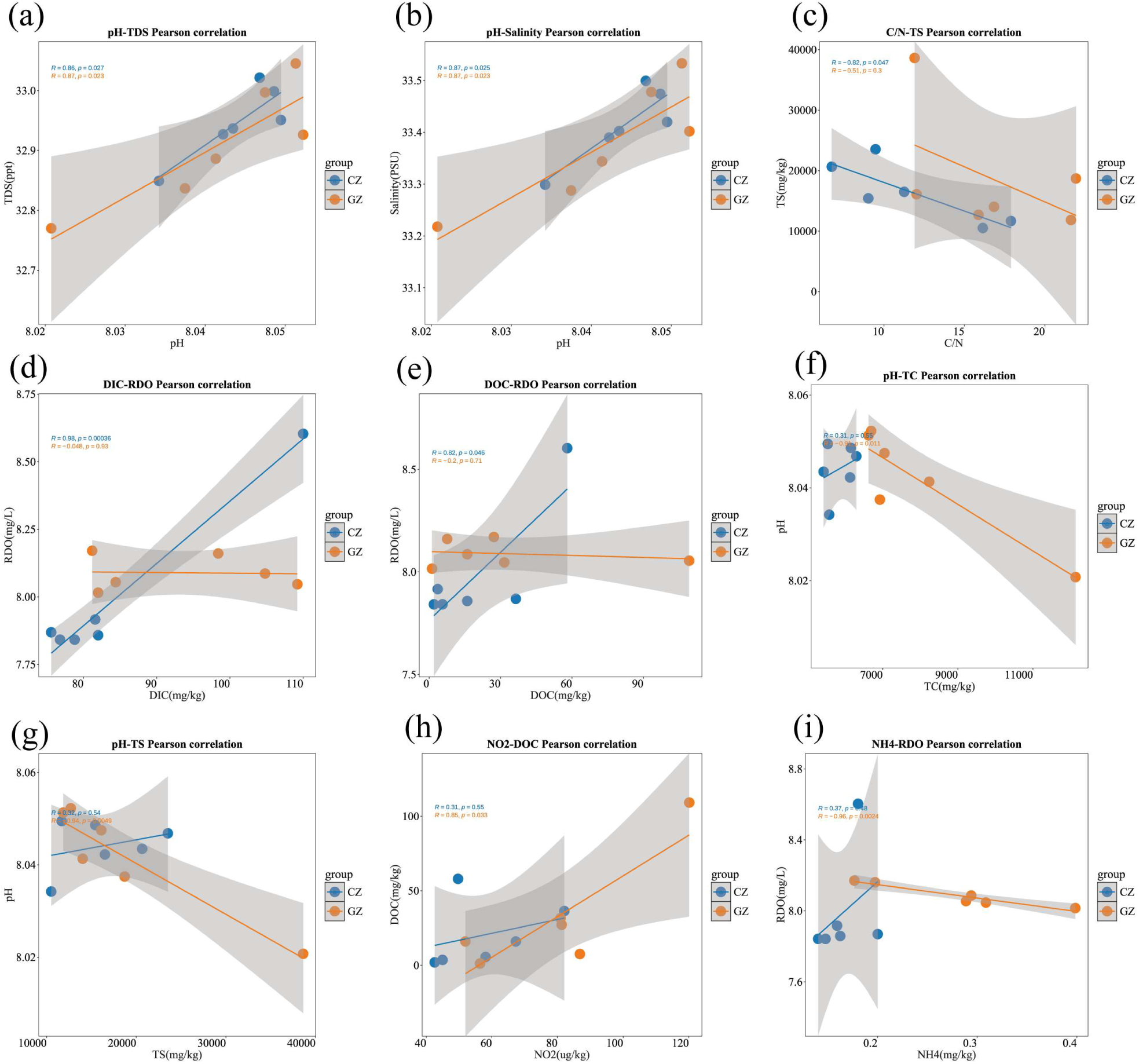
The linear correlations (Pearson’s) among physicochemical parameters of the sampling sites. (a) Overlying seawater TDS vs pH, (b) Overlying seawater salinity vs pH, (c) Sediment C/N vs TS, (d) Sediment DIC vs overlying seawater RDO, (e) Sediment DOC vs overlying seawater RDO, (f) Sediment TC vs overlying pH, (g) Sediment TS vs overlying pH, (h) Sediment NO_2_-N vs DOC and (i) Sediment NH_4_-N vs overlying RDO.

### 3.2 Microbial carbon fixation functional genes

After data screening and quality control, 11 target genes were detected **(Fig. 2, Tab. S2)**. Non-metric multidimensional scaling (n-MDS) based on Bray-Curtis distances revealed distinct clustering of carbon cycling gene compositions between the cultivation and control zones **(Fig. 2b)**, with significant structural differences confirmed by ANOSIM (*p* < 0.05; **Tab. S3**). The abundance of multiple carbon cycle-related functional genes was significantly higher in cultivation zones than in control zones (*p* < 0.05; **Fig. 2**). Specifically, carbon fixation gene abundance in cultivation zones was 5.21-fold higher than in control zone, with the Calvin-Benson-Bassham (CBB) cycle, 3-hydroxypropionate cycle, and reductive tricarboxylic acid (rTCA) cycle exhibiting significantly elevated activity (*p* < 0.05; **Fig. 2d**). The multifunctionality index for carbon fixation processes in cultivation zones was significantly higher than in control zones (Wilcoxon test, *p* < 0.05; **Fig. 2c**). The indices of ecosystem versatility were 0.8342 ± 0.3325 in cultivation zones and 0.4407 ± 0.4754 in control zones,, respectively.

**Figure 2.**
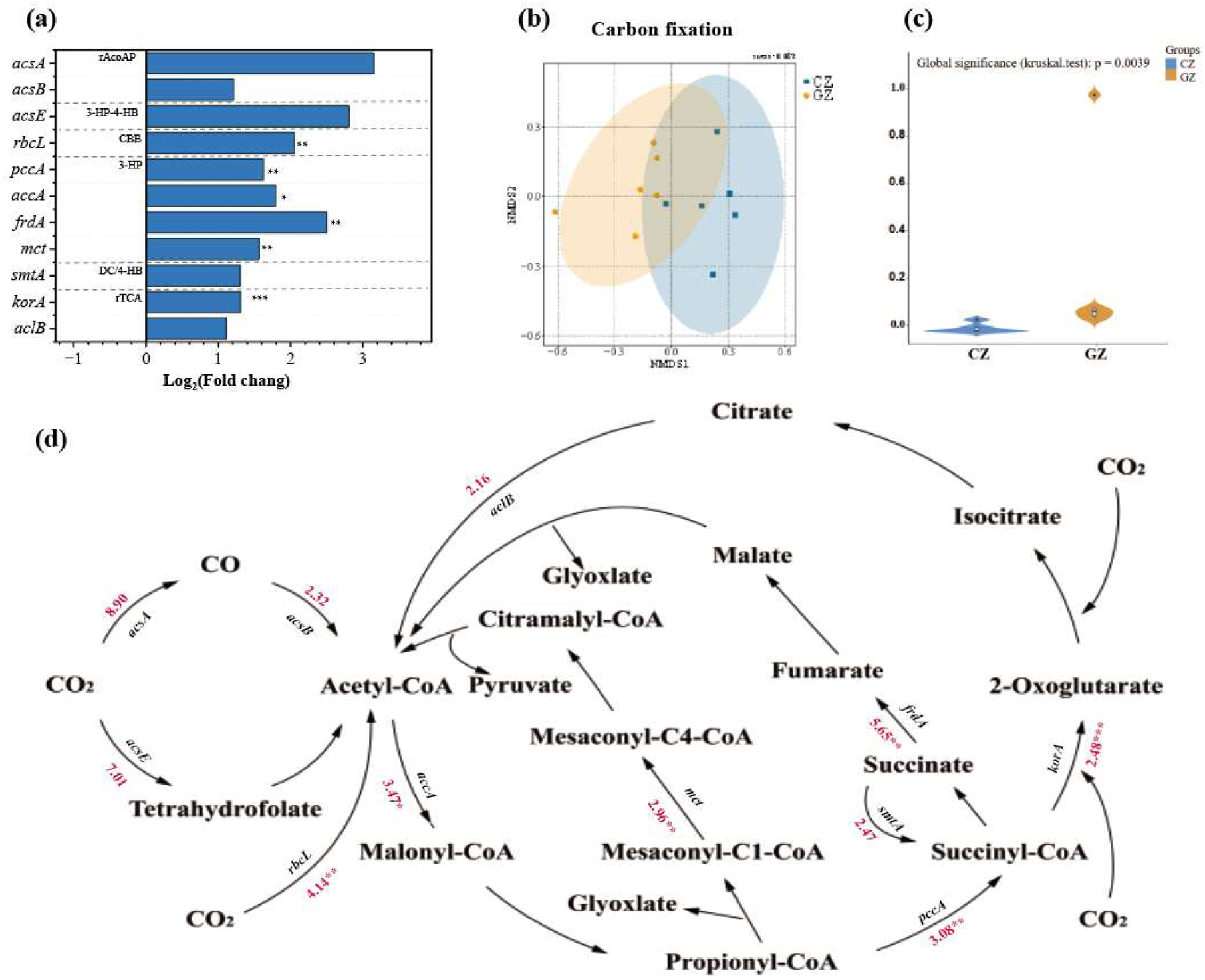
(a) Fold changes in the abundances of different carbon fixation genes between GZ and CZ samples. (b) NMDS analysis of carbon fixation functional genes. (c) Comparison of multifunctionality indices between GZ and CZ samples. (d) Carbon fixation pathways and the ratio of gene abundances between cultivation and control zones. Red values indicate the ratio of gene abundances in GZ relative to CZ samples. Significance was assessed using Student’s t-test: ***, *p* < 0.001; **, *p* < 0.01; *, *p* < 0.05.

### 3.3 Bacterial community diversity and composition

A total of 449,538 high-quality bacterial 16S rRNA sequences were obtained from sediment samples, yielding 12,316 operational taxonomic units (OTUs). Both richness (Chao1: 3,533–4,659 vs. 4,029–5,623) and Shannon diversity (6.557–6.998 vs. 6.804–7.399) were significantly lower in the cultivation zone than in the control zone **(Fig. 3a)**. In CZ, total carbon (TC) positively correlated with Chao1 and Shannon indices (one-way ANOVA, *p* < 0.05), whereas TC was inversely correlated with Chao1 in GZ. Total dissolved solids (TDS) positively correlated with both indices across combined (GZ + CZ) and zone-specific analyses **(Fig. S4)**.

**Figure 3.**
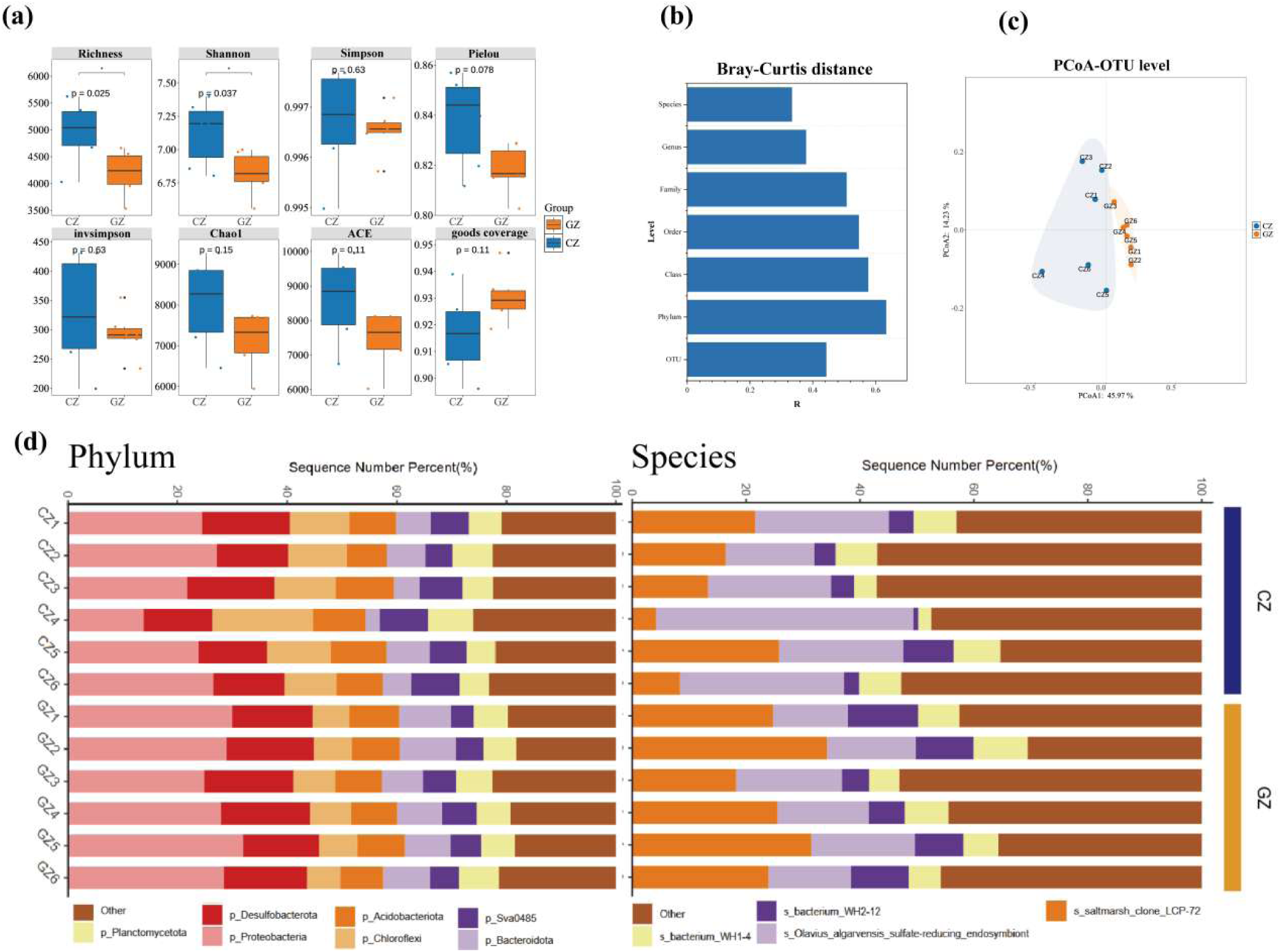
Bacterial community diversity and composition. (a) Alpha-diversity indices of the bacterial community. Significance was assessed using ANOVA with Tukey’s HSD test: *, *p* < 0.05. (b, c) Beta-diversity of bacterial communities based on Bray-Curtis distances, with multivariate homogeneity of group variances indicated. (d) Composition of dominant bacterial taxa at the phylum and species levels (relative abundance > 5%).

Bray-Curtis dissimilarity increased with taxonomic resolution **(Fig. 3b)**, and PERMANOVA confirmed significant differences in community structure between zones (*p* < 0.05). PCoA ordination (Bray-Curtis distance) revealed strong clustering of GZ bacterial communities **(Fig. 3c)**. At the phylum level, Proteobacteria, Desulfobacterota, Bacteroidota, Acidobacteriota, Chloroflexi, Planctomycetota, and Sva0485 dominated both zones, collectively accounting for 80% (GZ) and 77% (CZ) of relative abundance **(Fig. 3d)**. Dominant species included *saltmarsh clone LCP-72*, *Olavius algarvensis sulfate-reducing endosymbiont*, *bacterium WH2-12*, and *bacterium WH1-4*.

LEfSe analysis (LDA > 2) identified zone-specific biomarkers **(Fig. 4)**. GZ biomarkers included six phyla (e.g., Proteobacteria, Cyanobacteria) and nine species, such as *saltmarsh clone LCP-72*, *Fuerstia marisgermanicae*, and *delta-proteobacterium ectosymbiont* of *Rimicaris exoculata* (*p* < 0.05). CZ biomarkers comprised 18 phyla (e.g., Chloroflexi, Nitrospirota) and nine species, including *Olavius algarvensis sulfate-reducing endosymbiont* and *Truepera radiovictrix DSM 17093* (*p* < 0.05).

**Figure 4.**
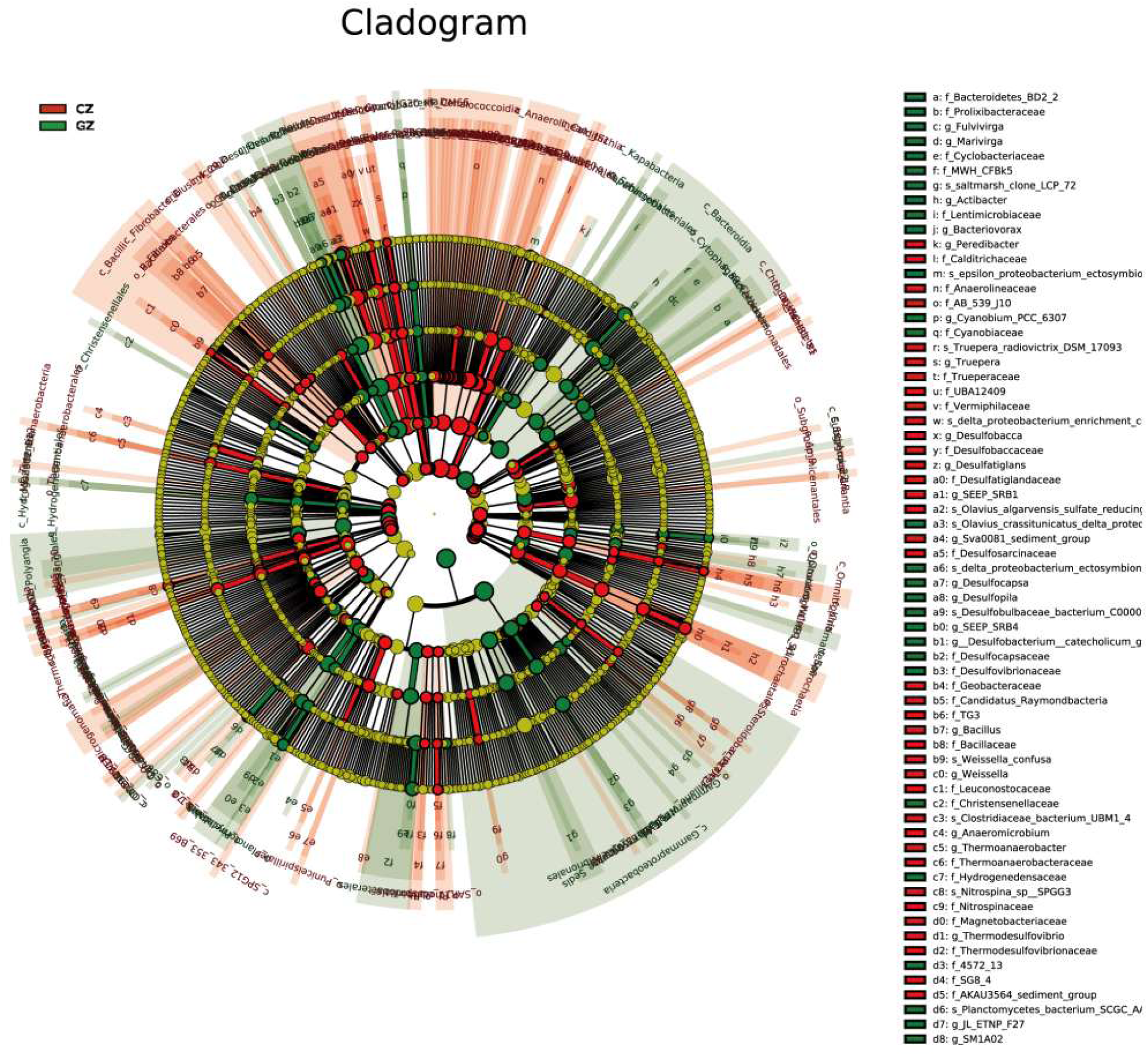
LEfSe (Linear Discriminant Analysis Effect Size) analysis of bacterial communities in cultivation (GZ) and control (CZ) zones. Cladogram depicting the bacterial community structure in GZ and CZ. Yellow nodes indicate taxa that did not differ significantly between the two zones. LDA score (log10) represents the linear discriminant analysis score.

### 3.4 Molecular ecological network of bacterial communities

The GZ network comprised 1,187 nodes and 1,145 edges, while the CZ network had 940 nodes and 2,068 edges. Network-level topological parameters, including average degree, average clustering coefficient, harmonic geodesic distance, and density, were higher in CZ than in GZ, whereas the R² of the power law and modularity were higher in GZ. The proportion of positive links was greater in CZ than in GZ **(Fig. S5)**. Subnetworks for GZ and CZ, constructed using the k-core algorithm, were used to identify key species with high connectivity **(Fig. 5b & 5d)**. The CZ subnetwork exhibited stronger connectivity, with 144 nodes and 740 edges compared to 144 nodes and 319 edges in GZ, indicating more bacterial connections in CZ.

**Figure 5.**
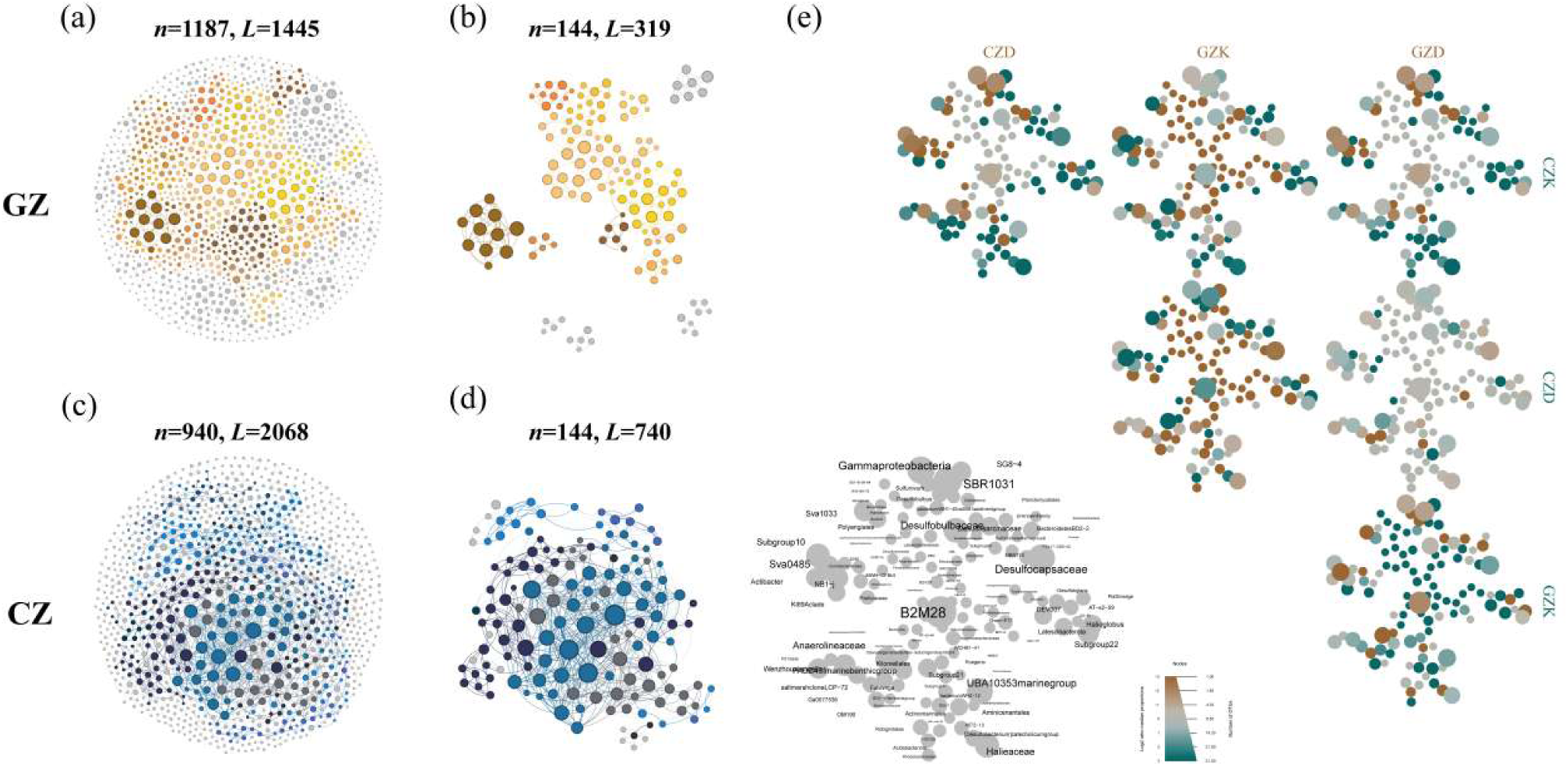
Molecular ecological networks of bacterial communities, k-core subnetworks, and the metacoder tree showing relationships between k-core and dominant species in GZ and CZ. (a, c) Molecular ecological networks of GZ and CZ, respectively. (b, d) K-core subnetworks of GZ and CZ, respectively. (e) Correlations among dominant species in GZ (GZD), dominant species in CZ (CZD), k-core species in GZ (GZK), and k-core species in CZ (CZK). In the networks and subnetworks, the top item indicates the number of nodes (n) and edges (L) in each network. Node size is proportional to degree (connectivity), and node color represents modules. In the metacoder tree, the lowest taxonomic rank of all species is displayed; node size reflects relative abundance, and node color indicates the strength of the association based on Spearman correlation coefficients.

The total number of species in GZ k-core (GZK), GZ dominant (GZD), CZ k-core (CZK), and CZ dominant (CZD) groups was 144, 88, 144, and 110, respectively. Most k-core species were not dominant (124 of 144 in CZK and 124 of 144 in GZK). The GZK and CZK groups exhibited significant differences, sharing only 18 species. A metacoder tree was constructed to explore correlations among GZD, CZD, GZK, and CZK species **(Fig. 5e)**. These analyses confirmed distinct species compositions between GZK and CZK, with GZK containing a higher proportion of low-abundance species, including members of *Latescibacteraceae*, *Rokubacteriales*, and *Hydrogenedensaceae*. Refer to **Fig. S6 & S7** for the specific species composition and differences of the four groups of bacteria.

**Table 1.**
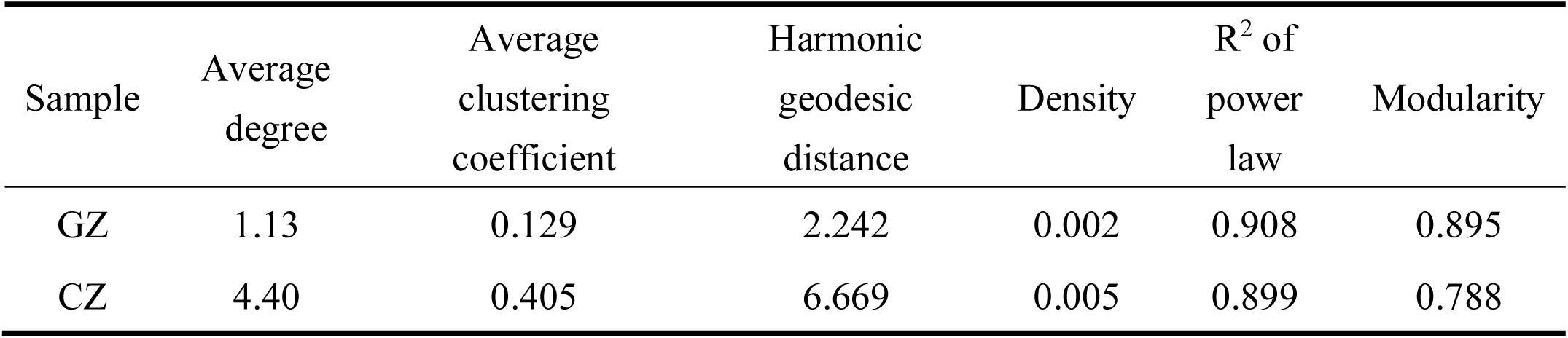
Topological properties of molecular ecological networks of bacterial communities in cultivation zone (GZ) and control zone (CZ)

### 3.5 Sediment carbon fixation rate

Rates exhibited significant spatial variation. In CZ, carbon fixation rates ranged from 6.44 × 10⁻⁸ to 1.40 × 10⁻⁶ g C h⁻² g⁻¹ (dry weight), whereas in GZ, rates ranged from 4.05 × 10⁻⁷ to 2.60 × 10⁻⁶ g C h⁻² g⁻¹ (dry weight). The sediment carbon fixation rate in GZ was significantly higher than in CZ (one-way ANOVA, *p* < 0.05; **Fig. S9**). In CZ, the highest rate was observed at site CZ4 and the lowest at site CZ3. In GZ, the maximum rate occurred at site GZ1 and the minimum at site GZ6. The detail data are presented in **Tab. S6**.

### 3.6 Relationships of carbon cycle gene abundance, sediment carbon fixation rates, and environmental parameter with dominant and k-core species

A significant positive correlation between the abundance of *mct* gene in sediments of aquaculture areas and the abundance of dominant bacterial species, but no similar correlation has been detected in natural sea areas **(Fig. 6a & 6c)**. Ammonia nitrogen was as a pivotal factors influencing the functionality of carbon fixation genes, exhibiting significant correlations with the abundance of mct, frdA, rbcL and korA in GZ (*r* > 0.9, *p* < 0.05, Spearman’s correlation; **Fig. 6e**). The abundance of *mct* gene was significantly related to the abundance of dominant species in aquaculture sediments. The abundance of OTU from the dominant species Acidobacteriota was significantly correlated with the abundance of mct gene, among which the species of Aminicenantales and Ilumatobacteraceae were significantly positively correlated **(Fig. 6e)**. Carbon fixation rates were significantly positively correlated with k-core species from Subgroup10 and MBNT15 (*r* > 0.9, *p* < 0.05, Spearman’s correlation; **Fig. 6e**). In CZ, sediment total nitrogen (TN) and the carbon-to-nitrogen (C/N) ratio were significantly positively correlated with both dominant and k-core species. In contrast, environmental factors in GZ showed no significant influence on these microbial communities **(Fig. 6b & 6d)**, suggesting abiotic factors play a key role in shaping microbial community structure in CZ.

**Figure 6.**
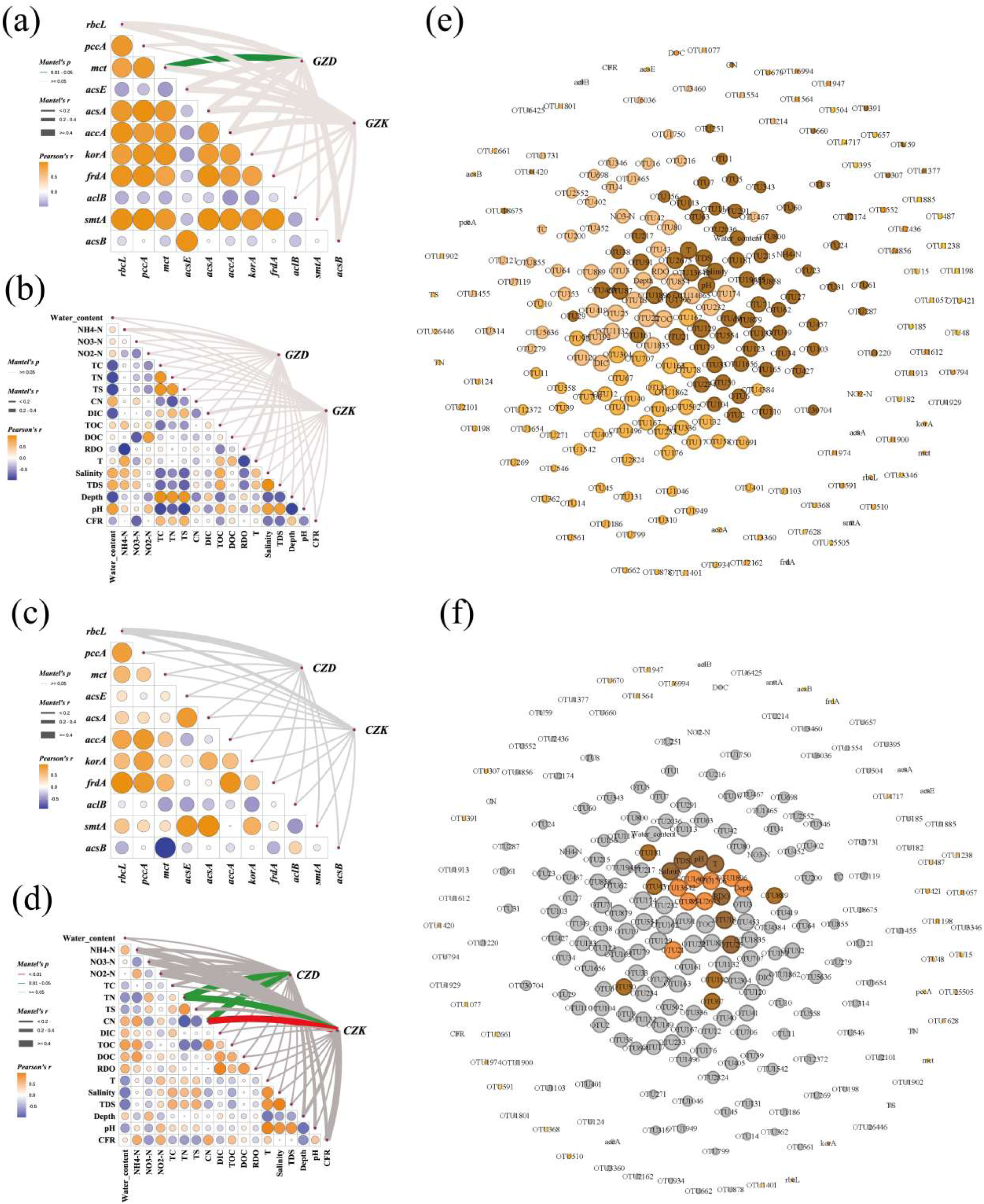
Relationships among sediment carbon fixation rates, carbon fixation gene abundance, environmental parameters, and dominant and k-core species. (a, c) Pairwise comparisons of carbon fixation gene abundances, with color gradients indicating Spearman’s correlation coefficients. K-core and dominant species were related to each environmental factor using partial Mantel tests based on Bray-Curtis distances. (b, d) Pairwise comparisons of environmental factors, with color gradients indicating Spearman’s correlation coefficients. K-core and dominant species were related to each environmental factor using partial Mantel tests. Edge width corresponds to the Mantel r statistic, and edge color indicates statistical significance: gray, *p* > 0.5; green, *p* = 0.01–0.05; red, *p* < 0.01. (e, f) Correlation network analysis showing relationships among sediment carbon fixation rates, carbon cycle gene abundance, and environmental parameters with dominant and k-core species in GZ and CZ. High correlations with significant edges (*r* > 0.9, *p* < 0.01, Spearman’s correlation) are displayed. Node size is proportional to relative abundance, and node and edge colors indicate network modules.

## 4 Discussion

### 4.1 Large-scale macroalage cultivation enhanced carbon fixation in coastal sediment

#### 4.1.1 Spatial heterogeneity of environmental factors in sediments

Interactions between environmental factors and microorganisms strongly influence ecosystem functions (García-Palacios et al., 2018; Xu et al., 2022; Sun et al., 2023). Large-scale macroalgal cultivation improves water quality by removing nutrients that drive eutrophication (e.g., NH₄-N, NO₃-N, NO₂-N, PO₄-P), increasing dissolved oxygen, and raising pH (Yang et al., 2006; Xu et al., 2008; Xie et al., 2017). In this study, concentrations of NH₄-N and total carbon were significantly higher in cultivation zone sediments than in control zones, while total organic carbon (TOC), dissolved organic carbon (DOC), and dissolved inorganic carbon (DIC) were slightly higher in GZ than in CZ (**Fig. S3; Tab. S1**), indicating that macroalgal cultivation increases sediment organic content (Bin et al., 2014). Macroalgal detritus contributes substantially to organic matter accumulation in sediments (Huang et al., 2024). In the control zone, overlying seawater relative dissolved oxygen (RDO) was significantly positively correlated with DIC and DOC, but no such correlations were observed in the cultivation zone (GZ) (**Fig. 1; Tab. S1**). Similar differences were observed for correlations between overlying seawater RDO and sediment NH₄-N, total carbon, and seawater pH versus sediment total sulfur. These variations likely result from microbial processes, macroalgal life activities, and hydrological changes induced by cultivation (Koomklang et al., 2018; Kang et al., 2024; Cui et al., 2021), suggesting that altered sediment–seawater interface processes drive differences in carbon cycling between cultivation and control zones.

#### 4.1.2 Spatial heterogeneity of carbon fixation genes in sediments

Coastal sediment ecosystems, particularly in large-scale macroalgal cultivation zones, exhibit high redox activity due to organic matter inputs, making them key areas for studying carbon cycling (Pessarrodona et al., 2022; Liu et al., 2022; Dyksma et al., 2016; Sun et al., 2023; Krause-Jensen et al., 2016; Buck-Wiese et al., 2023). Our results showed that large-scale macroalgal cultivation enhanced carbon cycling in coastal sediment ecosystems (**Fig. 2**). Carbon fixation was significantly higher in the cultivation zone than in the control zone. In particular, macroalgae cultivation significantly enhanced the activity of the 3-hydroxypropionate cycle, the reductive tricarboxylic acid (rTCA) cycle, and the Calvin cycle. Previous studies have demonstrated that these pathways are vital for carbon fixation within coastal sediment ecosystems and are often closely associated with chemoautotrophic microorganisms (Liu et al., 2022; Vineis et al., 2023). Although large-scale macroalgal cultivation is often considered a potential strategy for carbon neutrality (Gao et al., 2021; Wang et al., 2025), cultivation environments do not consistently act as CO₂ sinks due to the combined effects of macroalgae and microorganisms (Xiong et al., 2024). Surface sediments, ranging from a few millimeters to several centimeters in depth, have high dissolved oxygen concentrations and active redox reactions (Jørgensen et al., 2019). Organic matter accumulation in sediments can be transformed by microbial activity, increasing the release of nitrogen and inorganic carbon (Wang et al., 2022). Therefore, more direct evidence is needed to prove whether the breeding environment is a carbon source or a carbon sink.

#### 4.1.3 Spatial heterogeneity of multifunctional index and carbon fixation rate of sediments

While the potential of macroalgae to act as a carbon sink has attracted considerable attention, most studies have primarily focused on the direct carbon uptake by the macroalgae itself (Krause-Jensen et al., 2016; Gao et al., 2021; Wang et al., 2025). However, the role of microorganisms in carbon cycling should not be overlooked. Research in mangrove, estuarine, and other coastal ecosystems has shown that vegetation-rich habitats provide ample carbon sources that stimulate microbial carbon cycling (Mai et al., 2021; Hu et al., 2024; Liu et al., 2022). In macroalgae cultivation ecosystems, previous studies have mainly addressed microbial-mediated dynamics of particulate organic carbon (POC) and dissolved organic carbon (DOC), both of which contribute to the biological carbon pump (Lu et al., 2023; Li et al., 2022; Erlania et al., 2023; Huang et al., 2024; Xu et al., 2024). In the present study, we found that the multifunctionality index of the ecosystem—based on the abundance of carbon fixation genes—was significantly higher in cultivation zones compared to control zones (**Fig. 2c**), indicating enhanced microbial carbon fixation potential. Furthermore, ^13^C-labeled sodium bicarbonate was used as a sole carbon source to incubate sediments from the large-scale macroalgae cultivation ecosystem, allowing determination of sediment carbon fixation rates under laboratory conditions. Results showed that sediment carbon fixation rates were significantly higher in cultivation zones than in control zones (**Fig. S9**). Changes in the sediment–water interface processes in cultivation zones were an important factor driving this increase. Organic matter input from macroalgal cultivation resulted in significantly higher ammonia nitrogen and total carbon in sediments from cultivation zones compared to control zones (**Fig. S3**), promoting active redox reactions that drive chemoautotrophic carbon fixation (Howarth, 1984). These results highlight the carbon fixation potential of sediment microorganisms in aquaculture zones, although short-term incubation experiments may overestimate actual rates (Broek et al., 2018). Future studies should comprehensively assess the carbon sequestration capacity of large-scale seaweed cultivation across multiple scales and throughout the aquaculture cycle.

### 4.2 Large-scale macroalage cultivation change the microbial community: emphasize the differentiation of key species

#### 4.2.1 The utilization of organic matter shaped the community in the culture area

The ability of a microbial community to thrive in a given habitat depends on the alignment between community functionality and environmental conditions (Graham et al., 2016; Liu et al., 2020). In this study, we investigated bacterial community structure and species composition in sediments, and constructed molecular ecological networks. Significant taxonomic differences were observed between control sea communities and cultivation zone communities (**Fig. 3c**). Large-scale macroalgal cultivation strongly impacted the cultivation zone ecosystem, driving community differentiation (Xie et al., 2017; Xu et al., 2022; Xie et al., 2023; Lu et al., 2023; Xie et al., 2024). Bacterial communities in cultivation zone sediments exhibited lower species richness and simpler network structures compared to the control zone, consistent with previous findings from Nan’ao Island cultivation zones (Xie et al., 2017; Xie et al., 2024). In control zones, sediment total carbon (TC) was positively correlated with Chao1, Shannon, and ACE indices, while overlying water salinity and total dissolved solids (TDS) were significantly correlated with the Pielou index; however, these correlations were absent in cultivation zones. As a potent anthropogenic disturbance, aquaculture fundamentally alters sediment ecological dynamics. Deposition of algal detritus introduces substantial organic matter and nutrients into sediments, leading to significantly elevated TC and ammonium nitrogen and creating a eutrophic, homogenized sediment environment. Consequently, TC no longer acts as a limiting factor for microbial diversity, masking its natural correlation with diversity metrics. This dominant nutrient input also overrides the influence of other environmental parameters, such as salinity. Nutrient enrichment can promote microbial cooperation (Seo et al., 2009), resulting in simpler networks with fewer modules (Foster et al., 2012). Moreover, aquaculture activities may increase suspended solids in sediments, altering habitat conditions and microbial nutrient acquisition (Ramshaw et al., 2017; Wernberg et al., 2018). Dynamic changes in sediment pore water dissolved organic matter (DOM) directly affect bacterial community structure (Huang et al., 2024). In cultivation zones, DOM is primarily derived from microbial humic-like and tryptophan-like substances, whereas in control zones, DOM originates predominantly from terrestrial sources (Xu et al., 2022; Ou et al., 2024).

#### 4.2.2 Key species had the differentiation of community and potential carbon fixation function in culture area and control area

Key species are highly interconnected taxa within a microbiome that play a pivotal role in shaping community structure and function (Banerjee et al., 2018). This group includes both dominant and low-abundance species (Han et al., 2023; Sun et al., 2022). In this study, the K-core algorithm was employed to identify key species within the microbial networks of the cultivation (GZ) and control (CZ) zones. A metacoder tree was further used to differentiate dominant from key species across these zones (**Fig. 5**). Results showed that the majority of K-core species were not dominant species in either zone, with only 18 of the 144 K-core species shared between GZ and CZ (Fig. S6 & S7). Stable core species were observed across all four groups (GZD, GZK, CZD, CZK) at the phylum and class levels, particularly within Gammaproteobacteria, Desulfobacterota, and Bacteroidota, which were consistently abundant. These taxa are commonly reported in aquaculture sediments (Lu et al., 2023). K-core species in cultivation zones were significantly enriched with sulfate-reducing bacteria (SRB), including genera such as *Sulfurovum*, *Desulfobulbus*, and SBR1031. SRB are key players in the carbon cycle of anoxic marine sediments, contributing through anaerobic respiration, methanogenesis, and by providing electrons required for carbon fixation (Reyes et al., 2017; Ozuolmez et al., 2020; Stephania et al., 2021). Additionally, *Rhodobacterales* were prominent among K-core species in GZ. As facultative heterotrophs with carbon fixation potential, they interact closely with other microbial metabolisms, enhancing overall ecosystem carbon cycling (Kopejtka et al., 2017; Liu et al., 2021). Sediments from large-scale macroalgae aquaculture sites were also enriched with metabolically diverse microbes, particularly algae polysaccharide-degrading bacteria. Genera such as *Aquimarina*, *Fluviicola*, *Hirschia*, Bacteroidetes BD2-2, and *Psychroserpens* were notably abundant. Marine Bacteroidetes are known to degrade complex algal polymers (Krüger et al., 2019; Yu et al., 2024), and taxa like *Aquimarina* and the family *Flavobacteriaceae* have been reported as algal polysaccharide degraders (Lin et al., 2012; Zhou et al., 2015). Consequently, the microbial biomarkers identified in cultivation zone sediments formed a cohesive macroalgae-associated cluster, displaying both taxonomic and functional uniformity. The high abundance of algae polysaccharide-degrading microorganisms in these sediments likely facilitates enhanced carbon cycling in aquaculture sedimentary environments.

### 4.3 Linkages between carbon fixation and microbial community

#### 4.3.1 The availability of organic matter shapes the key species in aquaculture areas

Microorganisms play a pivotal role in enhancing sediment carbon fixation through multiple mechanisms, including the modulation of nutrient supply and resource allocation (Delgado-Baquerizo et al., 2016; Delgado-Baquerizo et al., 2017). Our results indicate that the availability of organic matter is a key driver of differences in bacterial-mediated carbon cycling between cultivation zones and natural marine environments. Specifically, total nitrogen (TN) and the carbon-to-nitrogen (C/N) ratio were significantly correlated with K-core species in natural marine sediments, whereas no such correlation was observed in cultivation zones (**Fig. 6b & 6d**). This pattern is consistent with observations from other coastal ecosystems, where organic matter quality and availability strongly influence microbial community structure and function (Hu et al., 2024; Delgado-Baquerizo et al., 2017). The absence of correlation in cultivation zones is likely due to the abundant and labile organic matter derived from aquaculture activities, which can be readily utilized by microorganisms. In contrast, surface sediments in natural marine environments are primarily composed of relatively recalcitrant terrestrial organic matter, imposing stronger constraints on microbial community composition and function (Sui et al., 2024; Huang et al., 2024).

#### 4.3.2 Functions of key species in carbon fixation cycle in aquaculture area

The abundance and composition of key microbial communities strongly influence the biogeochemical behavior of coastal sediments. In general, sulfur oxidation is the predominant chemoautotrophic process in reducing sediments, whereas ammonia oxidation plays a minor role (Howarth, 1984; Middelburg, 2011; Liu et al., 2022). However, in the aquaculture sediments of this study, ammonia nitrogen emerged as a pivotal factor affecting carbon fixation genes, showing significant correlations with the abundance of *mct*, *frdA*, *rbcL*, and *korA* in GZ (**Fig. 6e**). This may be due to the elevated ammonia concentrations in aquaculture sediments, which allow ammonia-oxidizing microorganisms to outcompete sulfur-oxidizers, enhancing their contribution to carbon fixation (Liu et al., 2022). The abundance of the *mct* gene, encoding monocarboxylate transporters, was strongly linked to dominant species in GZ, particularly OTUs from Acidobacteriota such as Aminicenantales and Ilumatobacteraceae. These taxa, often regarded as “microbial dark matter,” are prevalent in anaerobic, nutrient-rich environments and are capable of degrading complex organic matter, including aromatic and nitrogenous compounds (Dinh et al., 2022; Zakharova et al., 2022; Cheng et al., 2023; Silva-Solar et al., 2024). Their co-occurrence and high abundance in *Gracilariopsis* cultivation sediments suggest synergistic interactions in processing algal detritus and dissolved nutrients. Positive correlations between *mct* and these bacterial groups indicate a link between organic acid metabolism and carbon fixation. Furthermore, the correlation of *rbcL* (Calvin cycle) and *korA* (rTCA cycle) underscores the coexistence of multiple carbon fixation pathways, reflecting the metabolic versatility of aquaculture microbial communities—a phenomenon less commonly reported in coastal sediments (Berg et al., 2011; Chen et al., 2021; Jiang et al., 2022).

Consistent with gene abundance patterns, sediment carbon fixation rates in GZ were significantly higher than in CZ. K-core species analysis revealed strong positive correlations between carbon fixation rates and taxa such as Subgroup10 of Desulfobacterota and MBNT15 (**Fig. 6e**). Desulfobacterota are known to harbor carbon-fixing genes, particularly via acetate assimilation, contributing to coastal carbon sequestration (Dyksma et al., 2018; Yu et al., 2023). MBNT15, although typically rare in coastal sediments, was abundant here, suggesting a unique niche as a scavenger of low-molecular-weight organic compounds derived from microbial degradation of complex polymers (Begmatov et al., 2022). These results highlight the importance of active microbial oxidation reactions in providing terminal electron acceptors, thereby enhancing carbon fixation rates in surface sediments (Zhao et al., 2020; Liu et al., 2022). While our observations are limited to the macroalgal growth period, they provide a critical functional baseline for evaluating the carbon sink potential of large-scale macroalgae cultivation environments.

## 5 Conclusion

This study demonstrates that large-scale cultivation of *Gracilariopsis lemaneiformis* significantly enhances microbial-driven carbon fixation in coastal sediments. Cultivation markedly increased the abundance of key carbon fixation genes, particularly those involved in the Calvin-Benson-Bassham, 3-hydroxypropionate, and reductive tricarboxylic acid cycles. Correspondingly, sediment carbon fixation rates and the multifunctional index of the carbon fixation ecosystem were significantly elevated. These changes were accompanied by distinct shifts in microbial community structure, characterized by reduced diversity but enhanced functional specialization. Keystone taxa—including sulfate-reducing bacteria, dominant species of Acidobacteriota, and MBNT15—played pivotal roles in mediating carbon transformation and fixation. The simplified yet efficient microbial networks observed in cultivation zones further supported enhanced biogeochemical cycling. Overall, these findings highlight the potential of macroalgae cultivation as a nature-based strategy to strengthen coastal blue carbon sinks, offering valuable insights for sustainable aquaculture management and climate mitigation initiatives.

## Authorship contribution statement

P.B.P. and H.D. contributed to the study design. P.B.P. and Y.L.C. conducted experiments, bioinformatics analysis, data interpretation, and manuscript preparation/revisions. M.A., C.Y.W., W.H.Z. and H.D. helped with manuscript revisions. All authors read and approved the manuscript.

## Declaration of competing interest

The authors declare no competing interests.

## Ethics statement

Not applicable.

## Data access statement

The dataset(s) supporting the conclusions of this article is (are) included within the article and its additional file(s).

## Funding

This research was supported by several funding sources, including the earmarked fund for the China Agriculture Research System (CARS-50), 2025 Research on breeding technology of candidate species for Guangdong modern marine ranching (2025-MRB-00-001), and Fujian Province Marine Service and Fishery High-Quality Development Special Fund Project (FJHY-YYKJ-2024-1-18-6).

---

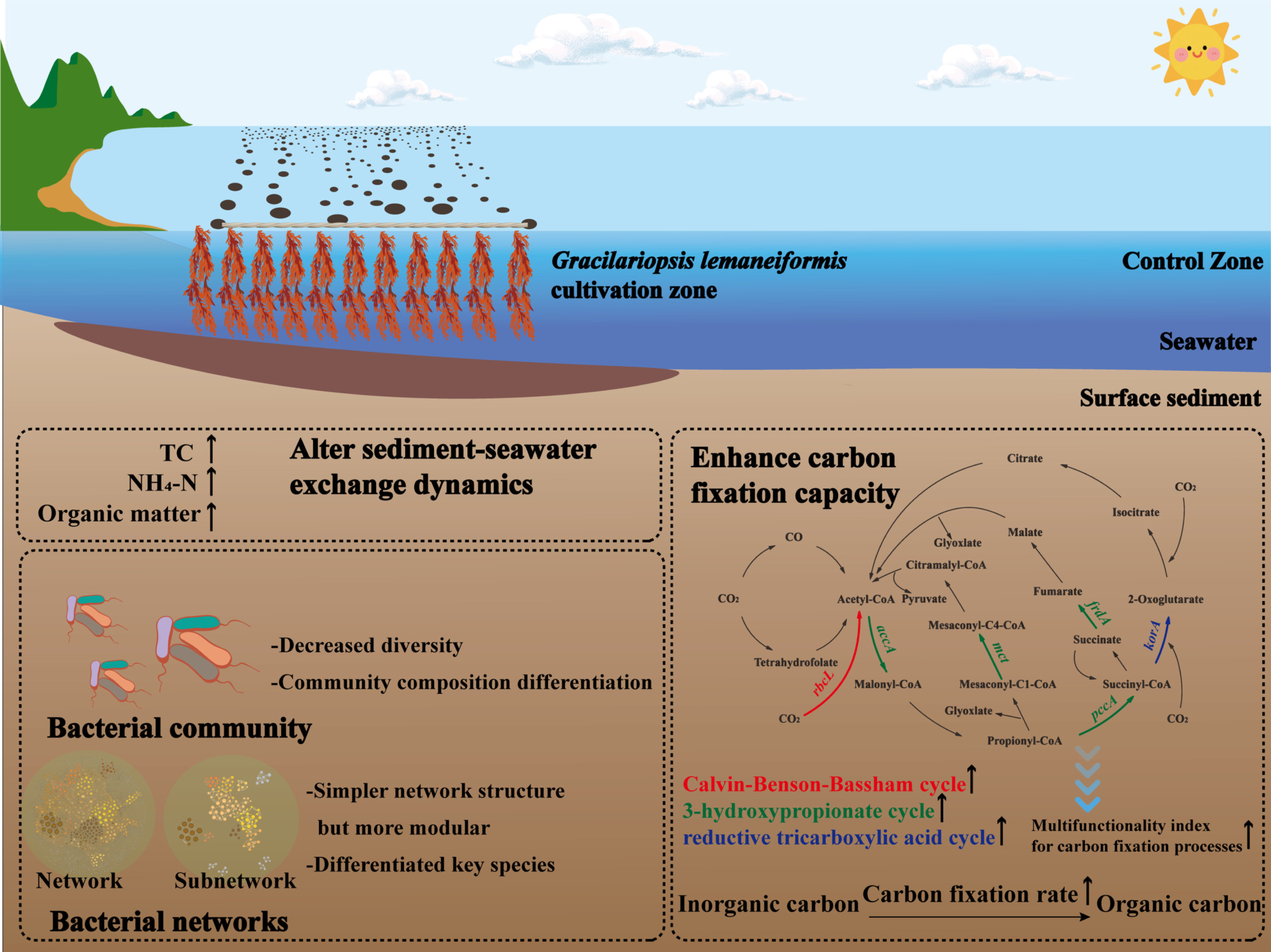

